# Evaluation of phenotypic and genotypic methods for the identification and characterisation of bacterial isolates recovered from catheter-associated urinary tract infections

**DOI:** 10.1101/2023.12.11.571106

**Authors:** Adam M. Varney, Eden Mannix-Fisher, Jonathan C. Thomas, Samantha McLean

## Abstract

**Purpose:** Urinary tract infections are the most common type of hospital-acquired infection, up to 80% of which are associated with catheterisation. The present study evaluates phenotypic and genomic characterisation of a panel of catheter associated urinary tract infection isolates from a UK hospital.

**Methods:** Strains were identified and characterised utilising a range of phenotypic and genomic techniques to understand where methodologies agree. The effect of medium composition on growth and biofilm formation phenotype was also determined to evidence the importance of assay design in characterisation of bacterial isolates.

**Results:** No consensus was observed for any of the CAUTI isolates across five identification methods, including biochemical testing, MALDI and sequencing technologies. Comparison of EUCAST antimicrobial susceptibility testing and genotypic data for antibiotic resistance showed high concordance where strains were phenotypically resistant to multiple antibiotic classes, however discordance increased for strains that were phenotypically sensitive to range of antibiotics. Phenotypic analysis of bacterial pathogens often relies on the use of rich laboratory media; however, we observed significant differences in growth rate and biofilm formation within a range of media, with a trend towards comparatively low planktonic growth and significant biofilm biomass formation in artificial urine.

**Conclusion:** This study emphasises potential pitfalls of relying on a single method of species identification, with only whole genome sequencing providing accurate identification of isolates to species level. Furthermore, it highlights the continuing importance of utilising phenotypic methods to understand antibiotic resistance within clinical settings and of utilising clinically relevant conditions for pathogen characterisation.

## Introduction

The Centers for Disease Control and Prevention recognise four top causes of hospital acquired infection, these include, catheter-associated urinary tract infection (CAUTI), central line-associated bloodstream infection, ventilator associated pneumonia and surgical site infection [1]. Within the UK healthcare setting, urinary tract infections account for 20-40% of all hospital associated infections [2], with 70-80% of those being catheter associated [3–6]. Catheters are therefore a major risk factor for infection and with 15-25% of all adult patients and up to 60% of intensive care patients having a urinary catheter placed at some stage during hospital admission the scale of the issue is significant [3,4,7,8]. The greatest risk factor for development of a catheter associated UTI is time, with the daily risk of infection ranging from 3-7% [3,9]. Patients with short term catheter placement (fewer than seven days of catheterisation) experience a 10-50% chance of infection rising to an almost 100% chance of infection in patients that are catheterised chronically for 28 days or more [10]. Although mortality rates for CAUTI are relatively low amongst hospital acquired infections (estimated at ∼2.3%), they are the leading cause of secondary nosocomial bloodstream infections; with approximately 17% of hospital acquired bacteraemia’s being of urinary origin, and thereby contributing to the mortality of ∼10% for this infection type [11].

Proportions of causative organisms in reported CAUTIs vary significantly between studies and geographical location [12]. The Centers for Disease Control and Prevention report that *Escherichia coli* cause around 21% of CAUTIs, with *Klebsiella pneumoniae* and *Pseudomonas aeruginosa* accounting for 7.7% and 10% respectively [8]. The CDC further report that *Candida* spp. account for 21% and *Enterococcus* spp. and *Enterobacter* spp. cause 14.9% and 4.1% of infections respectively [8]. The wide array of causative organisms is likely due to the catheter environment, where the catheter provides a scaffold for microbial adherence and proliferation prior to infection. Another factor is the varying health of patients during hospitalisation, with long-term catheterised patients often being intensive care patients that are more vulnerable to infection. Once microbes adhere to host cell surfaces or the catheter, their phenotype changes, allowing biofilm formation to occur through production of an extracellular matrix. This affords the microbial community within the biofilm additional protection from antimicrobials and host defences. Biofilms pose a significant risk to patient health with resistance to antibiotic treatment increasing up to 1,000-fold in this state. Biofilm formation also aids in evasion of the immune response creating resistance to phagocytosis and removal by physical forces. An additional complication of biofilm formation is the potential for physical blockage of the tubing that can lead to life-threatening urine retention, which backflows to the kidneys [13]. *Klebsiella* spp. and *Pseudomonas* spp. often cause blockage due to high mucoid production, although *Proteus* spp. have the greatest risk due to urease activity leading to crystalised biofilms that can become impossible to remove without surgery [14].

Identification of (CA)UTI is vital to wellbeing of the patient and knowledge of the isolate(s) present is required for successful treatment. Identification of (CA)UTI isolates is therefore undertaken in both clinical and research settings to aid in current treatment and to better understand the characteristics of pathogens to support future treatment options and drug / device development. Identification of these isolates varies between the two settings with research laboratories typically utilising a range of identification techniques including biochemical tests such as API, sequence-based approaches including 16S rRNA gene sequencing, or, when available, whole-genome sequencing with comparisons to a reference genome using Average Nucleotide Identity (ANI). In contrast the UK standards for microbiology investigation for examination of urine are followed by NHS clinical laboratories (SMI B 41: investigation of urine). However, the implementation of these standard procedures allows for local variations, particularly in the choice of culture techniques. Despite some variations, it is common practice for culture and sensitivity testing to be performed on positive urine cultures identified via flow cytometry. Isolates are initially cultured on CLED and / or chromogenic agar. CLED differentiates between lactose and non-lactose fermenters, whereas chromogenic agar allows for identification of common UTI pathogens at genus or species level depending on organism and the chromogenic agar of choice. Within the clinical setting the antibiotic profile of strains is often more critical than identification of isolates to species level and therefore species identification is not always necessary, for instance: Enterobacteriaceae (excluding *Salmonella spp*.) are often identified as “coliforms”, even though *Klebsiella spp*. and *E. coli* are distinguishable if using chromogenic agar; *Pseudomonas* spp. (commonly referred to as pseudomonads) are identified to genus level as are Enterococci.

Agar culture methods remain simple, cheap and the gold standard within the clinical setting, however, more advance techniques are available for identification including MALDI-TOF (Matrix-Assisted Laser Desorption/Ionization Time-of-Flight Mass Spectrometry) employing mass spectrometry or VITEK2 which uses biochemical testing and provides sensitivity testing. These methods, while faster, are more costly and often reserved for situations where rapid results are required, for example in at-risk groups such as patients in the ICU or patients with potential sepsis.

With three of the four major causes of healthcare associated infections involving medical devices and biofilm formation, there is an urgent need to understand microbial contamination of these materials. Studying biofilm formation on innate surfaces, such as catheters, requires experimental conditions that mimic the clinical environment as closely as practicably possible. *In vivo* studies provide conditions most closely related clinical infection; however, this is often unfeasible particularly in the early stages of research or drug development due to ethical concerns. In these circumstances *in vitro* testing is required, however there is significant variation in practice for these assay types, which can generate misleading data.

In this study we phenotypically and genotypically characterised seven clinically isolated (CA)UTI bacterial isolates to better understand the most appropriate techniques to use in clinical and research settings. We also evaluated biofilm formation in different media types and an *in vitro* Foley catheter contamination model to examine the effects of altering these experimental parameters on biofilm formation.

## Materials and methods

### Bacterial strains and culture conditions

Strains used in this study are listed in Table 1 and were obtained from the Nottingham University Hospitals Trust Pathogen Bank or the University of Sheffield [15]. Strains were typically plated onto Mueller-Hinton agar as required and liquid cultures grown in Mueller–Hinton broth, glucose defined minimal medium [16], artificial urine medium [17] or plasma-like medium [18,19] as required. For growth analysis; 100 µL of each desired medium was added to wells of a 96-well plate and overnight culture was added to the plates at a final concentration of 1%. Plates were incubated with shaking at 37°C in a BioTek Cytation 3 plate reader with turbidity measured at 600 nm every 15 min for 24 h.

**Table 1.** A summary comparison of phenotypic and genotypic identification of seven (CA)UTI bacterial isolates.

| Strain origin /designation | Species identification by MALDI | Score Value | 16S rRNA identification | Species identification by WGS | ANI (%) | Colour on UTI agar – presumptive identification | API 20E identification and code (% ID, T) |
| --- | --- | --- | --- | --- | --- | --- | --- |
| ST131 EC958 [15,36]<br>Human urine | N/A | N/A | *Escherichia fergusonii* / *Shigella flexneri* | *E. coli* | 98.34 | Pink – *E. coli* | 5104572<br>*E. coli* (99.5, 0.96) |
| NUH: 18Y000138<br>Urinary catheter | *Klebsiella pneumoniae* | 2.22 | *K. quasipneumoniae* | *K. quasipneumoniae* | 96.61 | Blue – *K. pneumoniae* | 1215773<br>*K. pneumoniae* (98.2, 0.93) |
| NUH: 18Y001710<br>Urinary catheter | *Klebsiella pneumoniae* | 2.26 | *K. quasipneumoniae* | *K. pneumoniae* | 98.88 | Blue – *K. pneumoniae* | 1205773<br>*Pantoea* spp. 2 (46.1, 0.95) |
| NUH: 18Y001733<br>Urinary catheter | *Enterobacter bugandensis* | 2.27 | *E. - quasiroggenkampii* / *roggenkampii* / *pseudoroggenkampii* / *quasimori* / | *E. roggenkampii* | 98.59 | Blue – *K. pneumoniae* | 3301563<br>*E. cloacae* (99.6, 0.59) |
| NUH: 19Y000094<br>Urinary catheter | *Enterobacter cloacae* | 2.30 | *E. hormaechei* | *E. hormaechei* subsp. *hoffmannii* | 99.03 | Pink - *E. coli* | 3305532<br>*E. cloacae* (99.2, 0.56) |
| NUH: 19Y000373<br>Urinary catheter | *Enterobacter cloacae* | 2.32 | *E. hormaechei* | *E. hormaechei* subsp. *xiangfangensis* | 95.69 | Blue – *K. pneumoniae* | 3205572<br>*E. cloacae* (92.6, 0.55) |
| NUH: 20Y000092<br>Urinary catheter | *Escherichia coli* | 2.47 | *E. fergusonii* | *E. coli* | 98.78 | None - unknown | 4144512<br>*E. coli* (69.5, 0.77) |

NUH: Nottingham University Hospitals Trust Pathogen Bank, Nottingham, UK. MALDI scores: 0.00-1.69: No organism identification possible, 1.70-1.99:
low-confidence identification, 2.00-3.00: high-confidence identification. \*As per Merck guidelines for UTI ChromoSelect Agar (16636). API 20E identification was performed using APIWEB™. 16S identified using Bakta annotated 16S genes via EZBioCloud.

### Phenotypic and biochemical identification of bacterial isolates

CAUTI were first identified by the NUH Pathogen Bank using an in-house MALDI Biotyper by comparison against reference library entries following identification standard method 1.1. For phenotypic identification, strains were plated onto UTI ChromoSelect agar (Merck) made as per manufacturer’s instructions and incubated overnight at 37°C. Strains were also identified using the API 20E kit commercially available from bioMérieux UK Ltd. (Basingstoke, UK) according to manufacturer’s instructions using freshly streaked bacterial cultures from Mueller-Hinton agar plates. Identification of strains was confirmed using APIWEB™.

### Disk diffusion assays

Disk diffusion assays were performed in accordance with the European Committee on Antimicrobial Susceptibility Testing (EUCAST) guidelines [20]. Bacterial suspensions diluted to McFarland turbidity standard of 0.5 were swabbed evenly across Mueller-Hinton agar plates. Disks were applied and plates were incubated for 18 h at 37°C prior to measuring the zone of inhibition (mm). Sensitivity and resistance were determined by comparison to EUCAST 2023 breakpoints [21]. EUCAST control strain ATCC 25922 was included to verify the results.

### 96-well microtiter biofilm assays

Overnight cultures were diluted to a turbidity of OD_600nm_ = 0.05 in fresh medium, which was then added in 100 µL amounts to the wells of 96-well plates and incubated statically at 37°C for 24 h. After incubation, microtiter plates were washed with 100 µL PBS twice. Biofilms were heat fixed at 60°C for 60 min prior to addition of 125 µL of 0.1% crystal violet to each well and incubation at room temperature for 20 min. Wells were washed three times with 150 µL phosphate buffered saline then inverted and left to dry overnight after which, 150 µL of acetic acid (33%) was added to each well with gentle shaking for 10 min before absorbance was measured at OD_540nm_ using a BioTek Cytation 3 plate reader.

### DNA extraction and whole genome sequencing

Genomic DNA was extracted from cell pellets using the GenElute™ Bacterial Genomic DNA kit (Merck) as per manufacturer’s instructions. DNA quality was checked using a NanoDrop microvolume spectrophotometer (ThermoFisher Scientific) and quantified using a Qubit 4 fluorometer (Invitrogen) high sensitivity dsDNA assay kit. A hybrid genome sequencing approach was used to generate data via Oxford Nanopore Technologies’ MinION and an Illumina MiSeq to achieve six closed genomes and one draft sequence.

The six CAUTI isolates were sequenced in-house using the Nextera XT Library Prep Kit and a V2 2×150 cycles reagent kit on the MiSeq platform. *E. coli* EC958 was sequenced on the Illumina NovaSeq6000 platform by MicrobesNG (Birmingham, UK), and combined with in-house Oxford Nanopore sequencing for genome assembly. Adapter sequences were trimmed from Illumina sequencing reads using Trimmomatic v0.39. Oxford Nanopore sequencing was completed on the Mk1B MinION. The native barcoding kit 24 V14 SQK-NBD114.24 was used for library preparation including multiplexing. Libraries were loaded onto a MinION R10.4 flow cell and run for 48 h. All software used for genome assembly and error correction were run using default parameters unless specifically stated. Oxford Nanopore raw fast5 data was basecalled using the super high accuracy (SUP) model of Guppy v6.5.7, and demultiplexed. Porechop v0.2.4 was used to trim adapter sequences, with middle- and end-threshold values of 85% and 95%, respectively. Filtlong v0.2.1 was used to trim reads by quality and by length, keeping reads with a minimum length of 1kb.

Overlapping Nanopore sequencing reads were *de novo* assembled using Flye v2.9.1. Genome assemblies were first polished using Nanopore reads, with four iterations of Racon v1.5.0 (with parameters -m 2 -g -2 -u), followed by Medaka v1.7.2. Assemblies were further polished using trimmed Illumina data with Polypolish v0.5.0, POLCA from the MaSuRCA v4.0.9 package and Nextpolish v1.4.1.

Whole genome sequences are available under BioProject: PRJNA1034027.

### Bioinformatic analysis

Genome completeness and contamination were assessed with CheckM v1.2.1 [22]. FastANI v1.32 was used to confirm the species of each CAUTI isolate, by calculating average nucleotide identity compared to type strains of the presumptive species’ genus (accession numbers available in supplementary material [23]). Bakta v1.8.1 (database v5.0) was used for annotating genes within genomes [24]. Bakta annotations associated with this article are available via figshare. Antimicrobial resistance markers were identified using Resistance Gene Identifier (RGI) v6.0.0 tool of the Comprehensive Antibiotic Resistance Database (CARD) v3.2.5 [[25]. Only resistance genes that showed a perfect or strict match with coverage for a given gene and that achieved ≥90% identity and read length in the database were included in this study. Phage elements were predicted using Phastest [26,27]. Virulence genes were identified using VFanalyser in the Virulence Factor Database (VFDB) [28]. WGS data was compared to the genus database for each strain, or to family databases where genus was not available.

16S rRNA gene sequences were extracted from Bakta annotations and each individual 16S rRNA gene copy was analysed through EZBioCloud and the top result taken. Where different copies provided different results, all were represented.

### Characterization of phenotypic and genomic concordance/discordance

Whole-genome sequence (WGS) data were compared with disk diffusion assay (DDA) data for all (CA)UTI isolates against 21 antibiotics (140 combinations). For each combination, concordance was the classification where WGS data were predicted to encode AMR genes for that class of antibiotic and the isolate had a phenotypic resistant profile (WGS-R/DDA-R) or WGS data were not predicted to encode AMR genes for that class of antibiotic and the isolate had a phenotypic susceptible profile (WGS-S/DDA-S) as described [29,30]. Discordance was the classification assigned where WGS data were predicted to encode AMR genes and the isolate had a phenotypic sensitive profile (WGS-R/DDA-S) or WGS data were not predicted to encode AMR genes and the isolate had a phenotypic resistant profile (WGS-S/DDA-R). WGS results were classified as “resistant” when one or several AMR genes were identified by CARD and allocated as the mechanism of resistance to that antimicrobial, and as “susceptible” when no resistance gene was identified after the CARD exclusion criteria were applied (described above).

### The Drip Flow Reactor® Foley catheter tubing contamination model

The Drip Flow Reactor® (DFR) was purchased from Biosurface Technologies Corporation with modification to allow tubing attachment [31,32] . 16F all-silicone Foley Catheters were purchased from BARD^®^. Catheter retention balloons were inflated approximately halfway, clamped and cut to approximately 6.5 cm in length. A 1/8^th^ female luer barb was attached to the cut end of the catheter prior to connection to the DFR chamber. The modified DFR was prepared to allow media flow into the chamber via the inflow port and drain through the Foley catheter by gravity after the chamber was filled to the eyelet. The DFR was sterilised by autoclave and artificial urine medium (AUM) filter sterilised. Once sterilised the media lines were connected, and inflow tubes connected to the peristaltic pump (Fig. S1).

Starter cultures were prepared by incubation at 37°C and 150 rpm shaking in 50 mL total volume of the appropriate medium from a 1% overnight culture. Upon reaching early-exponential growth phase, cultures were diluted to 10^5^ – 10^6^ CFU mL^-1^ in AUM and the tip of the Foley catheter piece was dipped into the diluted culture for 30 s. The contaminated Foley catheter was then immediately secured inside the DFR chamber and chamber sealed. The whole apparatus was transferred to 37°C and the flow rate of medium was set to 0.5 mL min^-1^ for the assay, to mimic urine output for catheterised patients [33,34]. The DFR was run for five days, after which samples were removed for quantitative and qualitative analysis.

### Scanning electron microscopy of catheter biofilms

Washed catheters were transferred to a petri dish and sliced into 1 mm thick sections. Of the sections, four were randomly selected and the biofilm fixed by sequential incubation in 4% paraformaldehyde (30 min), 50-100% ethanol (in 10% increments, 20 min each), 1:2 hexamethyldisilazane (HMDS)/100 % ethanol (20 min), 2:1 HMDS/100 % ethanol (20 min) and 100 % HMDS (overnight).

After fixing, each piece was sliced laterally, placed lumen side up and sputter coated with gold using a Quorum Q150R ES Sputter Coater creating a 5 nm layer of gold across the surface of the tube lumen. The samples were viewed, and secondary electron imaging was carried out using a scanning electron microscope (Jeol JSM-7100F LV FEG-SEM) with high vacuum at 15 kV.

## Results

### Identification of (CA)UTI isolates by MALDI, 16S rRNA sequencing, whole genome sequencing, UTI chromogenic agar, and API 20E testing shows varying agreement between methodologies

CAUTI isolates were initially identified by MALDI analysis, with one *Escherichia coli*, two *Klebsiella pneumoniae* and three *Enterobacter spp*. identified. 16S rRNA sequencing agreed with MALDI analysis to genus level, however there was significant disagreement at species level (Table 1). The genomes generated by short Illumina reads, and Oxford Nanopore long reads resulted in six closed sequences, with one strain sequenced by Illumina short reads as a draft sequence. Isolates were identified as *Escherichia coli*, *Klebsiella quasipneumoniae*, *Klebsiella pneumoniae, Enterobacter roggenkampii,* and *Enterobacter hormaechei* by ANI analysis against the type strain genome sequence for each respective species (>95-99% ANI [35], Table 1). This method represents the most comprehensive identification of strains within this study and provided further genotypic detail. WGS identified that isolates carried between one and six plasmids, ranging in size from 3632 - 274,807 bp. General features of the isolates are summarised in Table S1. Presence of prophage within genomes of the isolates were determined by Phastest (Table S2) and virulence factors identified via the Virulence Factor Database VFAnalyzer identified the presence of multiple virulence genes within the isolates, highlighting their pathogenic potential (Table S3).

Routine identification of (CA)UTI infection isolates in clinical settings often involves the use of UTI chromogenic agar. This agar contains various proprietary chromogenic substrates that when metabolised by common urinary tract infection strains produce different colours according to the metabolic pathways they possess, which can be utilised for identification. We sought to compare the presumptive chromogenic agar identification of the seven (CA)UTI isolates to their genomic identification to understand how reliable this common methodology is for this panel of isolates (Table 1). When the WGS identification was compared to presumptive identification on chromogenic agar; just two of the seven bacterial isolates tested had the expected chromogenic colour according to manufacturer guidance (*E. coli* EC958 and *K. pneumoniae). K. pneumoniae, K. quasipneumoniae* and two of the *Enterobacter spp*. as identified by WGS were blue suggesting the presence of *K. pneumoniae or Enterococcus faecalis,* which are readily distinguishable via colony size (*K. pneumoniae* being larger and more mucoid). Based on colony appearance on Chromogenic UTI agar the two *Enterobacter spp*. forming blue colonies would be classified as *K. pneumoniae* (Fig. S2). *E. hormaechei*^19Y000094^ showed pink colonies and would therefore likely be misidentified as *E. coli* during routine identification tests and *E. coli* ^20Y000092^ also displayed a non-conventional colour on chromogenic agar, with no colour change, which would also lead to potential misidentification (Table 1).

Another common testing method for bacterial identification is the use of biochemical testing such as those found in API testing kits. We identified the seven (CA)UTI isolates using the API 20E testing kit commercially available from bioMérieux UK Ltd. Identification via this method agreed with WGS identification to species level for the *E. coli* isolates only. API testing generally agreed with WGS identification to genus level, with the exception of *Klebsiella pneumoniae* being identified by API testing as *Pantoea spp*.

Comparison of identification methods mostly showed agreement to genus level for all methods except chromogenic UTI agar. At species level there was no clear agreement between identification methods with WGS required to reliably identify bacterial isolates (Table 1). All species names used hereafter refer to the species names as determined by whole genome sequencing as these identities are the most reliable.

### Genotypic and phenotypic antibiotic resistance profiles show varied concordance but do highlight the multidrug resistance commonly observed with (CA)UTI isolates

The EUCAST disk diffusion assay remains the gold standard method for antibiotic susceptibility testing to establish clinical resistance or sensitivity to many antibiotics for correct antibiotic treatment to be administered. The seven (CA)UTI isolates were tested for antibiotic susceptibility utilising EUCAST methodology [37]. Eleven unique classes of antibiotics and a total of 21 antibiotics were tested. These groups encompass the main treatment options for urinary tract infections set by the National Institute for Health and Care Excellence (NICE [38]) and included nitrofurantoin, trimethoprim and ampicillin (inferring amoxicillin sensitivity / resistance), which are the UK’s first line antibiotic treatments against urinary tract infections [38]. Cephalosporins from different generations, carbapenems, and fluroquinolones, the USA’s first line catheter-associated urinary tract infection antibiotic treatment prior to sensitivity testing, were also tested.

All strains except *E. coli^20Y000092^*were classified as multidrug resistant being resistant to at least one antibiotic in three separate classes [39]. The six MDR isolates were highly resistant, averaging resistance against 15 of the 21 antibiotics tested. *K. pneumoniae* was resistant to all tested antibiotics except gentamycin and tigecycline and *E. hormaechei^19Y000094^* was only sensitive to chloramphenicol and nitrofurantoin (Table 2). Due to the lack of EUCAST guidance on disk diffusion assay breakpoints, colistin was not tested, however resistance was subsequently inferred genotypically via the mechanisms of efflux and antibiotic target alteration (Table 3). Two of the seven strains showed resistance to nitrofurantoin, four to trimethoprim and six to ampicillin, highlighting the increasing emergence of resistance to first line antibiotic treatments within the UK. *E. coli^20Y000092^* showed sensitivity to all antibiotic classes, which is unusual for a CAUTI isolate in the current antimicrobial resistance climate (Table 2).

**Table 2.** All tested (CA)UTI bacterial isolates were multidrug resistant by phenotypic testing. (R) indicates resistance and (S) sensitivity in accordance with EUCAST guidelines (v13.1) [21]. For cephalosporins, number in parentheses indicate generation. N=3

| Group | Antibiotic | *E. coli*<br>EC958 | *K. quasipneumoniae*<br>18Y000138 | *K. pneumoniae*<br>18Y001710 | *E. roggenkampii*<br>18Y001733 | *E. hormaechei*<br>19Y000094 | *E. hormaechei*<br>19Y000373 | *E. coli*<br>20Y000092 |
| --- | --- | --- | --- | --- | --- | --- | --- | --- |
| Penams | Ampicillin | 0 (R) | 0 (R) | 0 (R) | 0 (R) | 0 (R) | 0 (R) | 17 (S) |
|  | Piperacillin | 7 (R) | 18 (R) | 0 (R) | 12.5 (R) | 0 (R) | 3.5 (R) | 22.5 (S) |
|  | Amox + Clav | 6 (R) | 21.5 (S) | 7 (R) | 3.5 (R) | 0 (R) | 0 (R) | 19.5 (S) |
| Cephamycins | Cefoxitin | 13 (R) | 0 (R) | 0 (R) | 0 (R) | 0 (R) | 0 (R) | 25 (S) |
| Cephalosporins | Cefixime (3<sup>rd</sup>) | 0 (R) | 22 (S) | 0 (R) | 0 (R) | 0 (R) | 0 (R) | 21 (S) |
|  | Ceftriaxone (3<sup>rd</sup>) | 13 (R) | 26 (S) | 0 (R) | 11.5 (R) | 0 (R) | 3.5 (R) | 28.5 (S) |
|  | Cefotaxime (3<sup>rd</sup>) | 0 (R) | 21.5 (S) | 0 (R) | 0 (R) | 0 (R) | 0 (R) | 26 (S) |
|  | Ceftazidime (3<sup>rd</sup>) | 11.5 (R) | 25 (S) | 0 (R) | 0 (R) | 0 (R) | 0 (R) | 24.5 (S) |
|  | Cefepime (4<sup>th</sup>) | 25.5 (R) | 27.5 (S) | 12.5 (R) | 29 (S) | 12 (R) | 20 (R) | 31.5 (S) |
| Carbapenems | Meropenem | 31 (S) | 24 (S) | 14 (R) | 23 (S) | 13 (R) | 25.5 (S) | 29.5 (S) |
|  | Doripenem | 30 (S) | 26 (S) | 14.5 (R) | 23.5 (R) | 14 (R) | 25 (S) | 28 (S) |
| Monobactams | Aztreonam | 20 (R) | 29 (S) | 9 (R) | 15 (R) | 23.5 (R) | 11 (R) | 29 (S) |
| Fluoroquinolones | Ciprofloxacin | 0 (R) | 22 (R) | 16.5 (R) | 26.5 (S) | 14 (R) | 21.5 (R) | 30.5 (S) |
|  | Norfloxacin | 0 (R) | 17.5 (R) | 15 (R) | 24 (S) | 9 (R) | 21 (R) | 26 (S) |
| Aminoglycosides | Amikacin | 15.5 (R) | 19.5 (S) | 14 (R) | 18 (S) | 0 (R) | 19 (S) | 20 (S) |
|  | Gentamicin | 17 (S) | 18.5 (S) | 17.5 (S) | 17 (S) | 0 (R) | 13 (R) | 18 (S) |
|  | Tobramycin | 7 (R) | 18 (S) | 8 (R) | 17 (S) | 0 (R) | 14 (R) | 16.5 (S) |
| Glycylcyclines | Tigecycline | 20 (S) | 15.5 (R) | 19.5 (S) | 17.5 (R) | 0 (R) | 19 (S) | 21 (S) |
| Diaminopyrimidines | Trimethoprim | 0 (R) | 17 (S) | 0 (R) | 23.5 (S) | 0 (R) | 0 (R) | 23 (S) |
| Phenicol | Chloramphenicol | 22.5 (S) | 15 (R) | 0 (R) | 17 (S) | 19 (S) | 0 (R) | 21.5 (S) |
| Nitrofurans | Nitrofurantoin | 20.5 (S) | 8 (R) | 10.5 (R) | 12 (S) | 14.5 (S) | 14.5 (S) | 16 (S) |

**Table 3.**
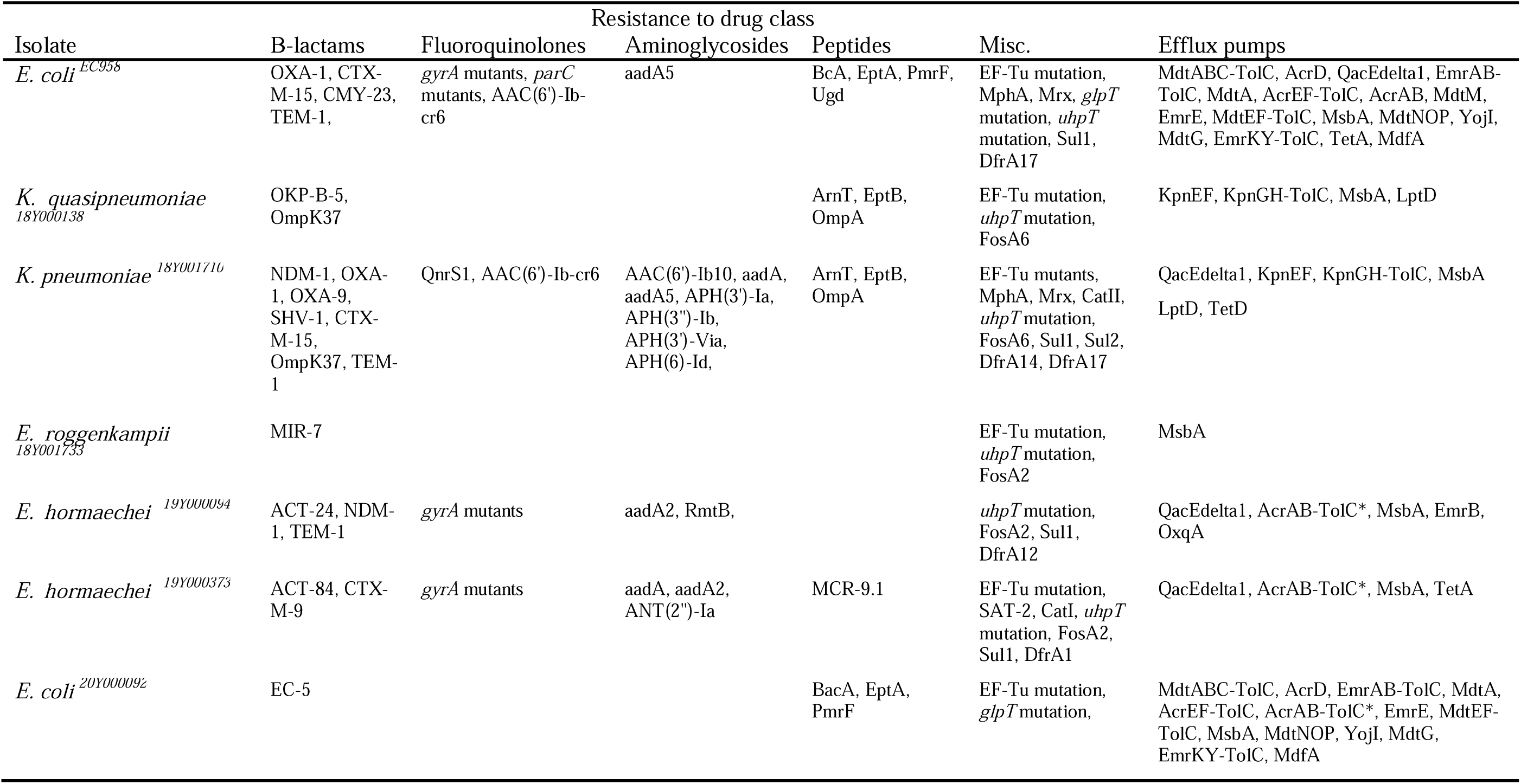
Predicted AMR genes encoded within genomes of the seven (CA)UTI isolates.

| Isolate | Resistance to drug class |  |  |  |  |  |
| --- | --- | --- | --- | --- | --- | --- |
|  | B-lactams | Fluoroquinolones | Aminoglycosides | Peptides | Misc. | Efflux pumps |
| *E. coli* <sup>EC958</sup> | OXA-1, CTX-M-15, CMY-23, TEM-1, | *gyrA* mutants, *parC* mutants, AAC(6')-Ib-cr6 | aadA5 | BcA, EptA, PmrF, Ugd | EF-Tu mutation, MphA, Mrx, *glpT* mutation, *uhpT* mutation, Sul1, DfrA17 | MdtABC-TolC, AcrD, QacEdelta1, EmrAB-TolC, MdtA, AcrEF-TolC, AcrAB, MdtM, EmrE, MdtEF-TolC, MsbA, MdtNOP, YojI, MdtG, EmrKY-TolC, TetA, MdfA |
| *K. quasipneumoniae* <sup>18Y000138</sup> | OKP-B-5, OmpK37 |  |  | ArnT, EptB, OmpA | EF-Tu mutation, *uhpT* mutation, FosA6 | KpnEF, KpnGH-TolC, MsbA, LptD |
| *K. pneumoniae* <sup>18Y001710</sup> | NDM-1, OXA-1, OXA-9, SHV-1, CTX-M-15, OmpK37, TEM-1 | QnrS1, AAC(6')-Ib-cr6 | AAC(6')-Ib10, aadA, aadA5, APH(3')-Ia, APH(3'')-Ib, APH(3')-Via, APH(6)-Id, | ArnT, EptB, OmpA | EF-Tu mutants, MphA, Mrx, CatII, *uhpT* mutation, FosA6, Sul1, Sul2, DfrA14, DfrA17 | QacEdelta1, KpnEF, KpnGH-TolC, MsbA, LptD, TetD |
| *E. roggenkampii* <sup>18Y001733</sup> | MIR-7 |  |  |  | EF-Tu mutation, *uhpT* mutation, FosA2 | MsbA |
| *E. hormaechei* <sup>19Y000094</sup> | ACT-24, NDM-1, TEM-1 | *gyrA* mutants | aadA2, RmtB, |  | *uhpT* mutation, FosA2, Sul1, DfrA12 | QacEdelta1, AcrAB-TolC*, MsbA, EmrB, OxqA |
| *E. hormaechei* <sup>19Y000373</sup> | ACT-84, CTX-M-9 | *gyrA* mutants | aadA, aadA2, ANT(2'')-Ia | MCR-9.1 | EF-Tu mutation, SAT-2, CatI, *uhpT* mutation, FosA2, Sul1, DfrA1 | QacEdelta1, AcrAB-TolC*, MsbA, TetA |
| *E. coli* <sup>20Y000092</sup> | EC-5 |  |  | BacA, EptA, PmrF | EF-Tu mutation, *glpT* mutation, | MdtABC-TolC, AcrD, EmrAB-TolC, MdtA, AcrEF-TolC, AcrAB-TolC*, EmrE, MdtEF-TolC, MsbA, MdtNOP, YojI, MdtG, EmrKY-TolC, MdfA |

Genotypic analysis of the seven (CA)UTI isolates using CARD predicted 113 antibiotic resistance genes present within the seven isolates, which is unsurprising due to the resistant nature of UTI pathogens and that the isolates are from three separate genera. Across the isolates, antibiotic resistance mechanisms included antibiotic efflux, antibiotic inactivation, target alteration, target replacement, target protection and reduced drug permeability. Multiple resistance genes for β-lactam antibiotics were predicted genotypically across the six resistant strains conferring resistance to penams, cephamycins, cephalosporins and carbapenems. Resistances to all major classes of antibiotic were predicted as well as generalised efflux mechanisms for resistance to antibiotics, disinfecting agents and antiseptics (Table 3). The greatest number of efflux pumps were found in the *E. coli* isolates, EC958 was phenotypically resistant to 15 of the 21 antibiotics tested, whereas *E. coli^20Y000092^*also contained a significant number of efflux pumps but was clinically sensitive to all antibiotics tested, suggesting that presence of efflux pumps is not a reliable indicator of phenotypic resistance.

To understand how the genotypic predicted resistance profiles of the seven strains compared to the phenotypic testing, we compared the two dataset and observed a concordance of 63.6%, with discordance at 36.4% within this panel (Fig. 1). Discordance was only observed when genotypic data identified one or more antibiotic resistances in an isolate genome, but phenotypic testing determined that the isolate was clinically sensitive to the antibiotic. Concordance also significantly varied between the isolates with *E. hormaechei^19Y000094^* and *K. pneumoniae^18Y001710^* showing concordance at 90%, whilst *E. coli^20Y000092^* showed 90% discordance. When the data was categorised by antibiotic; concordance varied between 28-85% with no obvious correlation between target antibiotic class nor mechanism of action (Fig. S3).

**Fig. 1.**
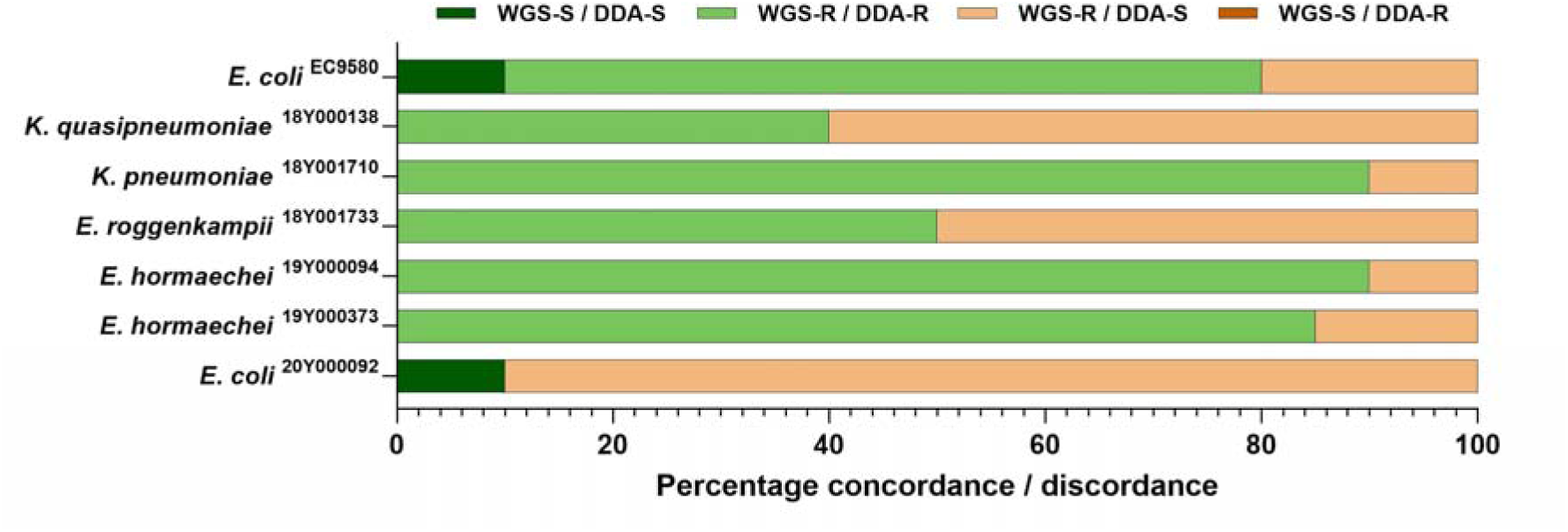
Genotype-phenotype percentage concordance and discordance for (CA)UTI isolate antimicrobial resistance

### Incubation of (CA)UTI strains in differing media conditions demonstrates varying metabolic capacity amongst the isolates

When characterising the phenotypic properties of bacterial isolates, common practice is to utilise a rich laboratory medium such as Mueller-Hinton broth. However, use of such medium can create a misleading phenotype for isolates that have evolved in conditions different to those of a rich laboratory medium. We therefore sought to determine the growth patterns of the seven (CA)UTI isolates in multiple media types to understand how they respond to different chemical environments. A rich medium (MHB) and defined minimal medium containing glucose (gDMM) were utilised to encompass commonly studied conditions and provide comparison to other reported research. Two physiologically relevant media were included in the study to understand how isolates grew in the presence of conditions that more closely mimic nutrient availability at different sites within the human body. Artificial urine medium (AUM) represented the environment encountered during (catheter-associated) urinary tract infection and plasma-like medium was used to understand how the strains responded to a medium containing components encountered in human plasma and interstitial fluid.

As expected, strains grew to the highest turbidity in the complex nutrient rich Mueller-Hinton broth. Where strains were able to grow in the defined minimal medium, they reached stationary phase quickly compared with the rich medium, likely due to depletion of an essential nutrient (Fig. 2). That two strains were unable to grow in the defined minimal medium is suggestive of a lower biosynthetic capacity compared with the other strains, possibly due to requirement for an essential nutrient present in urine but not within the minimal medium. For the two physiologically relevant media, the strains grew comparatively well in the plasma-like medium, but artificial urine medium had both a much lower final turbidity and lower maximum specific growth rate. Artificial urine medium is less nutrient rich than the other media types and contains urea, citric, boric and uric acids. Its pH is lower than other media at pH6.4, which could also contribute to lower growth rates and final cell densities.

**Fig. 2.**
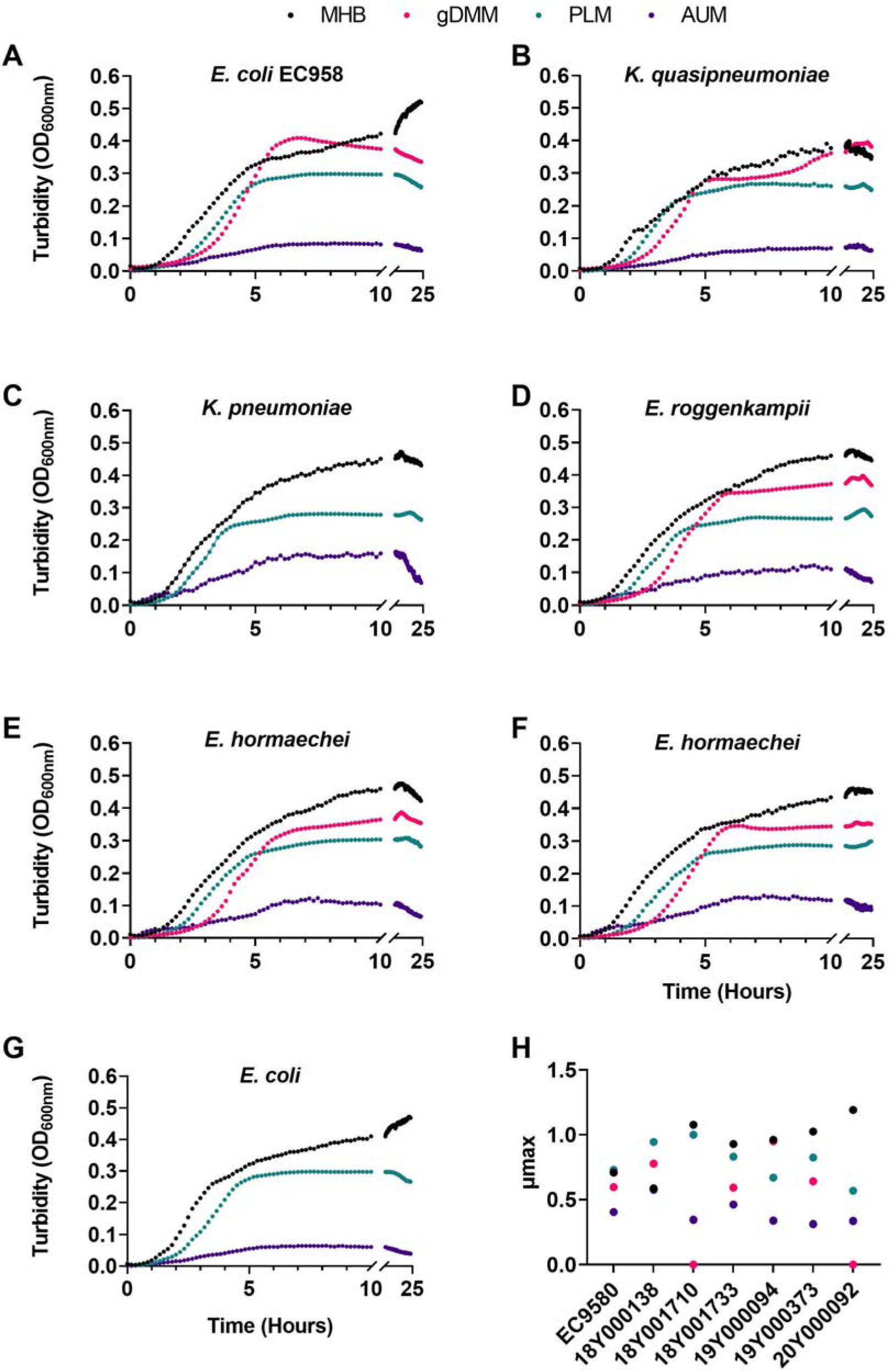
Growth of (CA)UTI bacterial isolates in different media show differing growth profiles Cultures were grown in 96-well plates from a 1% overnight culture inoculum from the same medium. Plates were incubated at 37 °C with shaking for 24 h with OD_600_ measured every 10 min. N = 4 with SD range 0.0004-0.1684. (A) *E. coli* ST131 EC958, (B) *K. quasipneumoniae*, (C) *K. pneumoniae^18Y001710^*, (D) *E. roggenkampii*, (E) *E. hormaechei^19Y000094^*, (F) *E. hormaechei^19Y000373^*, and (G) *E. coli^20Y000092^*. (H) Maximum specific growth rates were calculated for each isolate and medium combination.

### Biofilm formation is significantly altered in different media types and between different strains

(CA)UTI develop by forming biofilms within the catheter and urinary tract, therefore understanding the biofilm forming capabilities of this type of isolate is critical. The study of biofilms and clinical strains in general is most frequently conducted using high-throughput assays in nutrient-rich media that promote robust growth and strong biofilm formation. While this approach yields comparative data among clinical strains, it may not accurately replicate biofilms seen *in vivo* [40]. Although some studies do use relevant media particularly urine for UPEC studies [41], it remains uncommon practice perhaps due to either additional ethical considerations and / or cost. Artificial physiologically relevant medium can be used to bridge these approaches offering the advantage of both scalability and standardisation during product development by eliminating any batch-to-batch variation seen when using media derived from humans or animals directly. To understand how the (CA)UTI isolates form biofilms in different media types, we completed biofilm formation assays in the rich laboratory media Mueller-Hinton broth, Tryptic Soy broth and Lysogeny broth as these are the most used media in biofilm formation studies. We also tested biofilm formation in a defined minimal medium to compare biofilm formation when comparatively few nutrients are available and tested two physiologically relevant media: plasma-like medium and artificial urine medium to determine whether biofilm biomass was altered in media that more closely mimic the environment in which a (CA)UTI or other infection would occur.

Minimal biofilm biomass was established by any isolate in the define minimal medium (Fig. 3). Comparatively weak biofilm formation was observed with strains incubated in plasma-like medium except for the *Klebsiella* isolates that showed high relative biofilm formation. Comparison between the three rich media (MHB, TSB and LB) showed that biofilm formation across the isolates, was least consistent, but generally biofilm biomass was low across the strains tested. *K. pneumoniae* and *K. quasipneumoniae* provided the exception with relatively high levels of biofilm formation in MHB. Isolates incubated in LB and TSB showed more consistent biofilm formation, though *E. roggenkampii* and *E. coli* EC958 were very weak biofilm formers under these conditions.

**Fig. 3.**
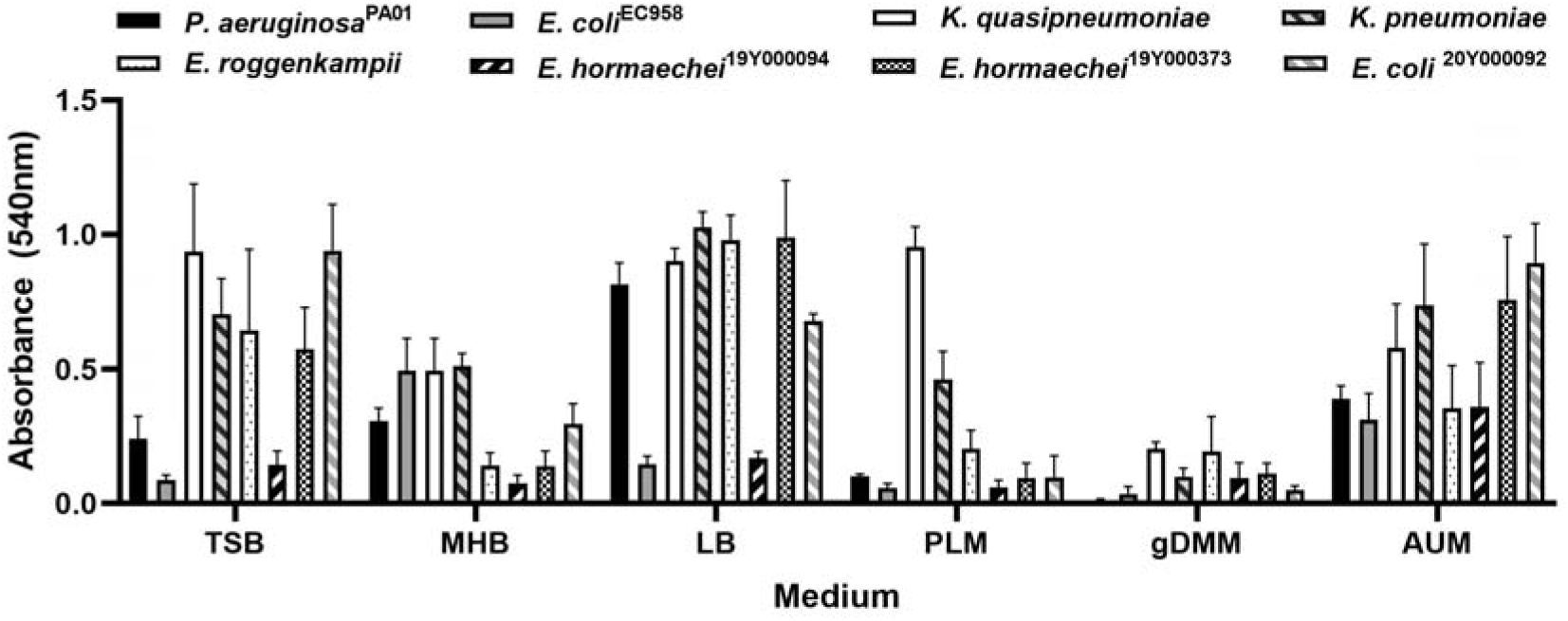
Biofilm formation varies significantly between strain and medium type Biofilms were grown statically in 96-well plates for 24 h at 37°C with an initial inoculation of OD_600_ = 0.05. *P. aeruginosa* PA01 was used as a positive control for biofilm formation. N = 3 ± SE.

Biofilm forming capacity of the (CA)UTI strains in AUM was varied. Comparison to the planktonic growth data (Fig. 2) highlights that whilst AUM supported the lowest growth rates and final turbidity, this does not hold true for biofilm formation, with AUM supporting comparatively high levels of biomass for several strains compared with MHB, gDMM and plasma-like medium (Fig. 3).

### Development of a clinically relevant biofilm model for assessment of CAUTI isolates

Whilst basic multi-well plate assays can be utilised for high throughput screening of biofilm formation capabilities and screening of antibiofilm drugs, the environment where the biofilm forms clinically is far removed from that of the multi-well plate assay. Therefore, we developed a Foley catheter bacterial contamination model to better mimic the *in vivo* conditions that lead to a CAUTI including flow, duration, catheter use, and medium composition. The Drip Flow Biofilm Reactor^®^ system was initially developed by Montana State University [42] utilising flat coupons to monitor biofilm formation on material surfaces. Bespoke modifications were developed by the manufacturer Biosurface Technologies Inc., that included fittings to allow tubing to act as the scaffold for biofilm formation. Utilising this modification to the Drip Flow Reactor^®^ we developed a system where the end of a Foley catheter replaced the flat material coupons. In this model, media flows into the DFR chamber, and is removed from the chamber by gravity flow through the islet of the Foley catheter, which better mimics the environment of a urinary catheter in a patient. In this work, the Foley catheters were contaminated by the bacterial isolate via dipping into a suspension of cells prior to attachment to the DFR chamber.

To better understand how CAUTI isolates form biofilms on urinary catheters, *K. pneumoniae^18Y001710^*and *E. hormaechei^19Y000094^* were selected for assessment in the Foley catheter contamination model. After five days incubation at 37°C, viable cells for the two isolates were cultured and were within the order of 10^10^ CFU mL^-1^. Live/Dead BacLight staining was used to determine the health of the biofilm, here the ratio of live to dead cells was within the range 40-60% (Fig. S4). Images of the biofilms formed in the microbial contamination model for both species were obtained by scanning electron microscopy, highlighting the structures formed in the urinary catheter contamination model for these species (Fig. 4).

**Fig. 4.**
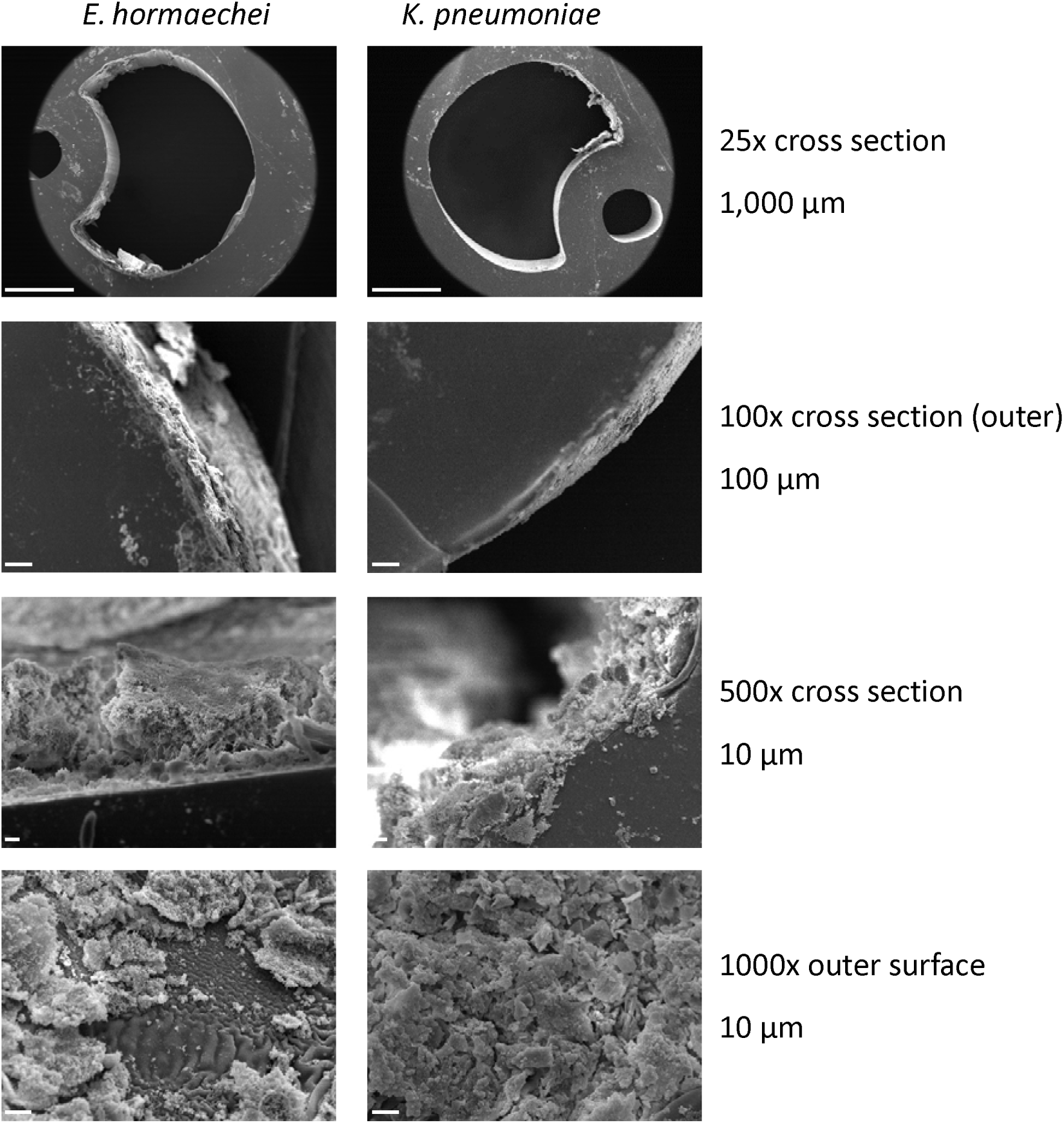
Scanning electron microscopy of CAUTI biofilms formed on Foley catheters in artificial urine medium in the Drip Flow Reactor^®^ Images show examples of biofilm growth at 15 kV on Foley catheters after five days incubation in artificial urine medium at 37°C.

## Discussion

Urinary tract infections account for around 40% of hospital acquired infections making them the leading cause of HAIs. With catheterisation in up to 80% of the UTIs reported in healthcare settings it is clear that catheterisation is a major risk factor for healthcare associated infections. This suggests that not only is it important to study the causative agents of these infections but also the manner in which they adhere to and proliferate on the catheter material prior to causing symptomatic bacteriuria. A hybrid WGS approach was undertaken to gain a deeper insight into seven (CA)UTI strain genotypes. Additionally, we identified isolates by 16S rRNA gene sequencing, MALDI and phenotypic means on UTI chromogenic agar and use of API 20 E testing to determine whether these approaches, common in clinical laboratories across the globe, provided consistent identification in agreement with sequencing.

In this study, plating onto UTI ChromoSelect agar classified *Enterobacter* isolates as *Klebsiella pneumoniae or E. coli* in accordance with the manufacturer’s instructions, furthermore *E. coli^20Y000092^*, showed no colour on the chromogenic agar making presumptive identification by this method impossible. The expected pink colour for this species is a result of β-D-galactosidase activity.

However, non-lactose fermenting and slow lactose fermenting *E. coli* have been shown to account for 10-20% of *E. coli* isolates [43,44], which are often more sensitive to cephalosporins, agreeing with antibiotic sensitivity testing (Table 2). Non-lactose fermenting *E. coli* strains are more likely to be community strains but this feature cannot be used as a predictive factor for infection as it does not affect the capability to cause disease compared with strains that are lactose fermenters [43,44]. This phenotype suggests that *E. coli^20Y000092^* is an opportunistic urinary pathogen, with WGS confirming the presence of numerous virulence factors but a lack of those that would be present in different pathovars of *E. coli* such as enteroaggregative or enteroinvasive (Table S3). *E. hormaechei^19Y000094^* colonies were pink when grown on UTI ChromoSelect agar. According to manufacturer guidance, the identities of all pink colonies should be confirmed by an indole test, since, some *Salmonella, Shigella, Citrobacter* and *Enterobacter* strains may also appear pink, which may be misidentified as *E. coli* but are indole negative, however this is at the discretion of the testing laboratory and may not always be undertaken to confirm [3]. Both *Klebsiella spp*. showed a typical blue colony appearance, and are readily distinguishable from *Enterococcus spp*. due to colony size. The lack of differentiation for *Enterobacter spp*. on chromogenic agar as well as relying on traditional biochemical markers highlights the dangers of relying on phenotypic testing to identify pathogens, with literature identification only correct to genus level 50% of the time [45].

Within the research literature, (CA)UTI causative organisms have been identified via multiple methods, however it is clear that appearance and chromogenic UTI profile is a key factor in identification as it is within the NHS, likely due to cost, leading to misrepresentation of species within the literature and clinical setting. Although, this is unlikely to affect treatment as sensitivity testing is also carried out prior to prescription of antibiotics where possible and critical infections are identified with additional methods, it could potentially skew medical literature altering the proposed significance of a strain within the clinical space [46]. This is potentially true with identification of *E. coli,* which within the literature is recorded to account for 75-90 % and 21% of all UTI and CAUTI respectively [47,48], however, if identification is based purely on UTI chromogenic agar, any Enterobacteriaceae with the correct biochemical profile could contribute to this statistic despite the likelihood of the true identity being a different genus.

Within this study, we also determined the antibiotic sensitivity of the strains both phenotypically and predicated genomically by CARD. The high levels of antibiotic resistance observed phenotypically across the (CA)UTI strains was unsurprising with the current state of global resistance. *K. pneumoniae^18Y001710^* was of particular concern being resistant to 19 of the 21 antibiotics tested, with potential colistin resistance mechanisms predicted via CARD (Table 3). Global resistance within *Klebsiella spp*. to colistin has been observed and is often a result of changes in cell permeability encoded on the chromosome. Strain NUH18Y001710 carries *arnT* and *eptB*, the products of which are involved in lipopolysaccharide modification that causes resistance to polymyxins [49][50–52]. *E. coli^20Y000092^* was atypical of CAUTI causing pathogens, being sensitive to all antibiotics tested.

However, antibiotic resistance genes identified from the genome sequence are suggestive of a resistance profile far greater than shown phenotypically. This raises interesting questions around the use of genotypic data to infer antibiotic resistance. Further investigation will be required to determine the cause of discordance between the genotypic and phenotypic AMR data for this isolate.

Sensitivity to one of the first line antibiotic treatments for UTI strains in the UK, nitrofurantoin, was observed in five of seven strains tested [38,53]. However, both *Klebsiella* isolates were resistant to this antibiotic, likely due to the predicted OqxA efflux pumps present in both *Klebsiella* isolates that are known to contribute to nitrofurantoin resistance [54].

As next generation sequencing is increasingly cost effective and readily available, and with new bioinformatics tools published that allow for basic genome annotation and interrogation with minimal bioinformatics expertise, it is tempting to utilise these technological advances to move to a more genotypic predictive system in place of phenotypic and biochemical testing. However, as demonstrated herein and elsewhere, it is clear that further development is required to reliably predict phenotype from genotype with a high degree of confidence [29,30,55]. Care should be taken when using genotypic tools to predict antimicrobial susceptibility. Using predictive tools alone could skew resistance data sets when comparing large numbers of strains genotypically and lead to incorrect treatment of a patient if used clinically without phenotypic testing. Although these tools are extremely useful for identification and tracking of resistance genes, phenotypic data should continue to be used to confirm the predictive tools, especially when susceptibility testing remains both fast and cost effective.

After identification, characterisation of isolates provides researchers with new avenues of investigation to prevent and / or treat infection. However, often these studies are performed within rich laboratory media containing many nutrients that the strains will not have access to during infection, or at significantly higher concentrations than found *in vivo*, which can alter bacterial metabolic processes and phenotype. Herein the (CA)UTI isolates were phenotypically characterised utilising various media to determine the effects of physiologically relevant chemical environments and improve understanding of how bacteria might respond during infection.

Growth curves and biofilm assays were both performed using a panel of media including rich laboratory media, defined minimal medium and two physiologically relevant media (AUM and PLM). Differences in media composition including between different types of rich medium resulted in differences in biofilm formation. Glucose often promotes biofilm formation [56,57] however, as the medium used in this study was a minimal medium, biofilm formation across the strains was low. Within *Staphylococcus* spp. TSB has been shown as a suitable medium for strong biofilm formation, more so than MHB [56], however MHB and TSB are comparably suitable biofilm formation media for *Pseudomonas* spp. [58] highlighting the differing responses of bacterial species to differing media components. MHB is a rich medium containing beef extract, hydrolysed casein, and starch, which can be broken down easily by bacteria to provide amino acids, carbon, vitamins, nitrogen, and other important nutrients. Therefore, it is not suitable for use in a clinically relevant CAUTI model as many of these nutrients will not be available, or available in the same concentrations, in human urine. In biofilm assays and models, researchers therefore often use these rich laboratory media at 50% of the recommended recipe strength to reduce the concentration of nutrients present, however this is still inconsistent with the chemical composition found in a clinical environment. AUM is much closer to the chemical composition of human urine and is able to promote strong biofilm formation for species including *P. aeruginosa*, particularly as part of a mixed biofilm [59,60]. Human urine would theoretically be the most appropriate medium for characterisation of urinary pathogens, however it was not used in this study for multiple reasons; firstly it requires ethical approval and incurs significant cost to acquire, particularly where larger volumes are required as in the Drip Flow Reactor model. Secondly, patient urine will vary greatly with factors including host gender, age, diet and disease conditions such as cancer or pregnancy. Even pooled urine suffers from batch to batch variation based on donor profile [61]. Therefore, to support standardisation and reproducibility of data in a medium that better mimics the infection environment, AUM is an appropriate medium that replicates *in vivo* conditions in a consistent manner [62].

The data present within this study underscores the need for caution when identifying strains to species level without the use of whole genome sequencing. It further highlights the need to study CAUTI isolated strains in clinically relevant conditions to more accurately understand how these pathogens will act *in vivo*, which would better support drug development in early pre-clinical phases. The data presented also exemplifies the antimicrobial resistance crisis, demonstrating on a small scale the significant resistance emerging in pathogens isolated within healthcare settings with all but one strain herein classified as multidrug resistant. Overall, this work demonstrates the need for a combination of phenotypic testing and genomic analysis to accurately identify and characterise clinically isolated bacterial pathogens so that we may overcome this threat to global health.

## Acknowledgements

The authors would like to thank Nottingham Trent University for funding via Vice Chancellors Awards and an Early Career Fellowship awarded to Dr Samantha McLean.

## Author contributions

All authors contributed to the study conception and design. Material preparation, data collection and analysis were performed by all authors. The first draft of the manuscript was written by Samantha McLean and all authors commented on previous versions of the manuscript. All authors read and approved the final manuscript.

## Data availability

The genomic data generated during the current study are available in the National Center for Biotechnology Information BioProject: PRJNA1034027. BioSample accession numbers: SAMN38049403-9. SRR raw Illumina data: SRR26595501-7. SRR raw Nanopore data: SRR26624650-5. Annotated genomes are available in <u>figshare</u>.

