## Supplementary data for "Evaluation of phenotypic and genotypic methods for the identification and characterisation of bacterial isolates recovered from catheter-associated urinary tract infections"

### Methods

#### Catheter biofilm viability assays

To carry out viability and live/dead staining tests of the biofilms, catheters were recovered from the DFR and washed with 3 ml sterile PBS. The catheter was then transferred to a fresh tube and biofilm removed with repeated cycles of pipetting, vortexing, and sonication in an Ultrawave Q series sonicating machine with 10 mL sterile PBS until no biofilm was observed by visual inspection. Viable counts were performed to calculate the number of live culturable cells. Viability was also determined by use of the LIVE/DEAD® BacLight™ Bacterial Viability Kit (Invitrogen) according to manufacturer's instructions.

**Table S1** Summary information for genomes generated from the (CA)UTI isolates described in this study

| Strain origin /designation | Species identification | Isolated | Coverage (x) | Completeness (%) | Contamination (%) | Length (bp) | Contigs | GC (%) | N50 (bp) | CDS |
|---|---|---|---|---|---|---|---|---|---|---|
| ST131 EC958 [19,37] Human urine | *Escherichia coli* | 03/2005 | 187 | 99.96% | 0.11% | 5,249,404 | 3 | 50.8 | 5,109,722 | 4,905 |
| NUH: 18Y000138 Urinary catheter | *Klebsiella quasipneumoniae* | 11/12/2015 | 124 | 99.99% | 0.6% | 5,262,360 | 1 | 58.1 | 5,262,360 | 4,830 |
| NUH: 18Y001710 Urinary catheter | *Klebsiella pneumoniae* | 09/05/2017 | 83 | 100% | 0.48% | 5,829,258 | 7 | 56.7 | 5,243,929 | 5,450 |
| NUH: 18Y001733 Urinary catheter | *Enterobacter roggenkampii* | 13/06/2017 | 138 | 99.96% | 0.45% | 4,658,510 | 2 | 56.1 | 4,617,805 | 4,216 |
| NUH: 19Y000094 Urinary catheter | *Enterobacter hormaechei* | 08/01/2019 | 29 | 99.78% | 0.19% | 4,805,056 | 155 | 55.1 | 58,874 | 4,513 |
| NUH: 19Y000373 Urinary catheter | *Enterobacter hormaechei* | 23/07/2019 | 124 | 99.73% | 0.19% | 5,203,737 | 3 | 54.7 | 4,769,210 | 4,895 |
| NUH: 20Y000092 Urinary catheter | *Escherichia coli* | 14/03/2019 | 100 | 99.96% | 0.08% | 4,846,350 | 2 | 50.6 | 4,733,682 | 4,525 |

NUH: Nottingham University Hospitals Trust pathogen bank

**Table S2** Predicted prophages present in the isolated genomes using PHASTEST

| Strain | Completeness | Score | Region Length | GC % | # Total Proteins | Most Common Phage |
|---|---|---|---|---|---|---|
| *E. coli* EC958 | questionable | 80 | 20.3Kb | 46.30% | 23 | PHAGE_Escher_SH2026Stx1_NC_049919(3) |
| | intact | 150 | 56.1Kb | 50.32% | 62 | PHAGE_Entero_BP_4795_NC_004813(9) |
| | intact | 150 | 43.4Kb | 48.06% | 60 | PHAGE_Pectob_ZF40_NC_019522(12) |
| | intact | 150 | 53.5Kb | 51.93% | 67 | PHAGE_Entero_BP_4795_NC_004813(25) |
| | intact | 150 | 44.8Kb | 51.09% | 52 | PHAGE_Entero_DE3_NC_042057(22) |
| | intact | 150 | 63.5Kb | 52.76% | 50 | PHAGE_Burkho_phiE255_NC_009237(28) |
| | intact | 150 | 50.4Kb | 52.07% | 58 | PHAGE_Entero_P88_NC_026014(40) |
| *K. pneumoniae* 18Y000138 | intact | 150 | 45.4Kb | 52.86% | 64 | PHAGE_Erwini_vB_EhrS_59_NC_048198(13) |
| | intact | 150 | 66.5Kb | 51.70% | 60 | PHAGE_Salmon_SPN3UB_NC_019545(11) |
| | intact | 150 | 38.6Kb | 53.45% | 31 | PHAGE_Klebsi_phiKO2_NC_005857(20) |
| | questionable | 90 | 40.1Kb | 54.82% | 20 | PHAGE_Klebsi_ST13_OXA48phi12.1_NC_049453(5) |
| *K. pneumoniae* 18Y001710 | questionable | 75 | 16.7Kb | 50.29% | 13 | PHAGE_Entero_P4_NC_001609(7) |
| | questionable | 75 | 51.4Kb | 55.49% | 25 | PHAGE_Escher_500465_1_NC_049342(12) |
| | intact | 150 | 44.5Kb | 55.14% | 41 | PHAGE_Erwini_EtG_NC_047833(28) |
| | intact | 150 | 56.1Kb | 50.11% | 51 | PHAGE_Klebsi_phiKO2_NC_005857(20) |
| | intact | 150 | 71.6Kb | 52.21% | 75 | PHAGE_Salmon_118970_sal3_NC_031940(16) |
| | intact | 140 | 40.3Kb | 50.11% | 35 | PHAGE_Salmon_SJ46_NC_031129(4) |
| | incomplete | 60 | 35.3Kb | 52.71% | 40 | PHAGE_Escher_RCS47_NC_042128(4) |
| *E. cloacae* 19Y000094 | incomplete | 40 | 20.2Kb | 50.40% | 15 | PHAGE_Bacter_APSE_2_NC_011551(5) |
| | intact | 150 | 41.4Kb | 51.92% | 53 | PHAGE_Entero_mEp390_NC_019721(22) |
| | intact | 150 | 57.4Kb | 52.71% | 91 | PHAGE_Salmon_SPN3UB_NC_019545(17) |
| | intact | 150 | 58.3Kb | 50.50% | 83 | PHAGE_Klebsi_phiKO2_NC_005857(24) |
| | intact | 150 | 33.2Kb | 52.11% | 49 | PHAGE_Escher_HK639_NC_016158(13) |
| *E. cloacae* 19Y000373 | incomplete | 20 | 21.2Kb | 49.34% | 22 | PHAGE_Entero_mEp237_NC_019704(10) |
| | questionable | 75 | 26.3Kb | 52.92% | 23 | PHAGE_Escher_500465_1_NC_049342(12) |
| | incomplete | 50 | 28.2Kb | 50.69% | 20 | PHAGE_Entero_P4_NC_001609(6) |
| *E. coli* 20Y000092 | intact | 120 | 45.1Kb | 52.15% | 59 | PHAGE_Escher_HK639_NC_016158(11) |
| | intact | 150 | 45.8Kb | 51.08% | 62 | PHAGE_Entero_mEp390_NC_019721(20) |
| | intact | 120 | 45.7Kb | 49.51% | 56 | PHAGE_Escher_phiV10_NC_007804(34) |
| | intact | 116 | 14.1Kb | 41.03% | 20 | PHAGE_Entero_I2_2_NC_001332(7) |
| | intact | 150 | 69.8Kb | 49.14% | 80 | PHAGE_Entero_DE3_NC_042057(21) |
| | intact | 150 | 54.4Kb | 50.22% | 61 | PHAGE_Entero_SfI_NC_027339(29) |
| | intact | 150 | 36.7Kb | 49.87% | 47 | PHAGE_Klebsi_4LV2017_NC_047818(28) |

Intact (score > 90) Questionable (score 70-90) Incomplete (score < 70). Orange text indicates prophage intact present on a plasmid and purple text indicates incomplete prophage on a second plasmid. See PHASTEST website for score classification criteria [26,27]. *Enterobacter asburiae* had no prophage regions predicated by Phastest.

**Table S3A** Virulence factors predicted by VFanalyzer using the Virulence Factor Database (Adherence)

| Isolate ↓ | Virulence Factor Class - Adherence | | | | | |
|---|---|---|---|---|---|---|
| | Virulence factors | | | | | |
| | *E. coli* common pilus (ECP) | P fimbriae | Type I fimbriae | Type III fibriae | Curli fibers | *Misc.* |
| *E. coli* [EC958] | *ecpA, cpB, ecpC, ecpD, ecpE, ecpR* | *papI* | *fimA, fimC, fimD, fimE, fimF, fimG, fimH, fimI* | | | AAF/II fimbriae: *aafC.* EaeH: *eaeH.* Lateral flagella: *flgC* |
| *K. quasipneumoniae* | | | *fimA, fimB, fimC, fimD, fimE, fimF, fimG, fimH, fimI, fimK* | *mrkA, mrkB, mrkC, mrkD, mrkF, mrkH, mrkI, mrkJ* | | |
| *K. pneumoniae* | | | *fimA, fimB, fimC, fimD, fimE, fimF, fimG, fimH, fimI, fimK* | *mrkA, mrkB, mrkC, mrkD, mrkF, mrkH, mrkI, mrkJ* | | |
| *E. roggenkampii* | | *papC* | *fimD, fimA, fimC* | *mrkC* | *csgA, csgD, csgF, csgB, csgC* | ELF – *elfC.* Streptococcal plasmin receptor – *plr/gapA*. pH6 antigen – *psaC.* CFA/I fimbriae: *cfaB* |
| *E. hormaechei* [19Y000094] | *ecpC* | *papC* | *fimA, fimC, fimD* | | *csgA, csgE* | pH6 antigen:, *hofB, hofC.* HCP pilus: *hcpA, hcpB, hcpC* |
| *E. hormaechei* [19Y000373] | *ecpC* | *papC* | *fimA, fimC, fimD* | *mrkC,* | *csgA, csgD, csgE, csgF, csgG* | pH6 antigen – *psaC* |
| *E. coli* [20Y000092] | *ecpA, ecpB, ecpC, ecpD, ecpE, ecpR* | | *fimA, fimB, fimC, fimD, fimE, fimF, fimG, fimH, fimI* | | | EaeH: *eaeH* |

**Table S3B** Virulence factors predicted by VFanalyzer using the Virulence Factor Database (Iron Uptake)

| Isolate ↓ | Virulence Factor Class – Iron uptake | | | | | | |
|---|---|---|---|---|---|---|---|
| | Aerobactin siderophore | Heme uptake/transport | Iron/manganese transport | Virulence factors Yersiniabactin siderophore | Ent siderophore | Salmochelin | Enterobactin transport |
| *E. coli* EC958 | *iucA, iucB, iucC, iucD, iutA* | *chuA, chuS, chuT, chuU, chuW, chuX, chuY* | *sitA, sitB, sitC, sitD* | *fyuA, irp1, irp2, ybtA, ybtE, ybtP, ybtQ, ybtS, ybtT, ybtU, ybtX* | | | |
| *K. quasipneumoniae* 18Y000138 | *iutA* | | | | *entA, entB, entC, entD, entE, entF, entS, fepA, fepB, fepC, fepD, fepG, fes* | *iroE, iroN* | |
| *K. pneumoniae* 18Y001710 | *iutA* | | | *fyuA, irp1, irp2, ybtA, ybtE, ybtP, ybtQ, ybtS, ybtT, ybtU, ybtX* | *entA, entB, entC, entD, entE, entF, entS, fepA, fepB, fepC, fepD, fepG, fes* | *iroE, iroN* | |
| *E. roggenkampii* 18Y001733 | *iucA, iucB, iucC, iucD, iutA* | *shuA, shuS, shuV, chuA, chuS, chuU, pvdH, viuC* | | | *entA, entB, entC, entE, entF, entS, fepA, fepB, fepC, fepD, fepG, fes* | | *fepA, fepB, fepC, fepD, fepG* |
| *E. hormaechei* 19Y000094 | *iucA, iucB, iucC, iucD, iutA* | *chuA, chuS, chuU, shuA, shuS, shuV, pvdH, viuC* | *sitA, sitB, sitC, sitD* | | *fepC, entA, entB, entC, entE, entF, fepA, fepB, fepC, fepD, fepG, fes* | *iroN* | |
| *E. hormaechei* 19Y000373 | *iucA, iucB, iucC, iucD, iutA* | *chuA, chuS, chuU, shuA, shuS, shuV, pvdH* | *sitA, sitB, sitC, sitD* | | *entA, entB, entC, entE, entF, entS, fepA, fepB, fepC, fepD, fepG,* | *iroN, orf04057* | |
| *E. coli* 20Y000092 | | *chuA, chuS, chuT, chuU, chuW, chuX, chuY* | *sitA, sitB, sitC, sitD* | *fyuA, irp1, irp2, ybtA, ybtE, ybtP, ybtQ, ybtS, ybtT, ybtU, ybtX* | | | |

**Table S3C** Virulence factors predicted by VFanalyzer using the Virulence Factor Database (Secretion System)

| Isolate ↓ | Virulence Factor Class – Secretion System | | | |
|---|---|---|---|---|
| | Virulence factors | | | |
| | SCI-I T6SS | T6SS | T2SS | T3SS |
| *E. coli* EC958 | *orf02153, orf02152, orf02151, orf02150, orf02149, orf02148, orf02147, orf02139, orf02142, orf02146, orf04638, orf04636, orf04635, orf04637, orf02131, orf02128, orf02127, orf02125, orf02124* | | | |
| *K. quasipneumoniae* | | *orf00728, orf00729* | | |
| *K. pneumoniae* | | *orf03749, clpV/tssH, dotU/tssL, hcp/tssD, ompA, vasE/tssK, vgrG/tssI, vipA/tssB, vipB/tssC, orf02845, orf02873, icmF, impA, impF, impG, impH, impJ, sciN* | | |
| *E. roggenkampii* | *orf02922, orf02942, orf02941, orf02940, orf02937, orf02936, orf02933, orf02930, orf02929, orf02928, orf02927, orf02923* | *orf02933, clpV/tssH, hcp/tssD, impA/tssA, vgrG/tssI, vipA/tssB, vipB/tssC, orf02942, orf02937, orf02936, orf02930, vasA/impG, orf02927, orf02928, impA, impF, orf02923, orf02921, clpV1, orf02424, orf02423* | *exeD, exeF, exeG, gspE, gspG, yst1E, yst1G, epsE* | *iagB* |
| *E. hormaechei* 19Y000094 | *orf01769, orf01768, orf01767, orf01766, orf01765, orf01764, orf03901, orf01763, orf01762, orf03943, orf03944, orf03945, orf03947, impA, impF* | *clpV/tssH, dotU/tssL, hcp/tssD, icmF/tssM, impA/tssA, ompA, sciN/tssJ, tssF, tssG, vasE/tssK, vgrG/tssI, vipA/tssB, vipB/tssC, clpV, clpV1, yst1O, exeD, exeF* | *epsE, yst1O, exeD, exeF, exeG, gspE, gspG* | |
| *E. hormaechei* 19Y000373 | *orf03537; orf03986, orf03538; orf03985, orf03539; orf03984, orf03540, orf03542, orf00721; orf02792; orf03543; orf03981, orf03544; orf03980, orf03546; orf03979, orf03551, orf03552; orf03971, orf03553; orf03970, orf03554; orf03969, orf03555, orf03556; orf03967* | *clpV/tssH, dotU/tssL hcp/tssD, icmF/tssM, impA/tssA, ompA, sciN/tssJ, tssF, tssG, vasE/tssK, vgrG/tssI, vipA/tssB, vipB/tssC, impF, impG, impH, impJ, orf04161, orf04160, clpV1* | *Orf03096, yst1E, yst1G, yst1O, exeD, exeF, exeG, gspE, gspG, epsE* | *iagB* |
| *E. coli* 20Y000092 | | *orf02463, orf02445, aec14, aec15, aec16, aec17, aec18, aec19, aec22, aec23, aec24, aec25, aec26, aec27/clpV, aec28, aec29, aec30, aec31, aec32, aec7* | | |

**Table S3D** Virulence factors predicted by VFanalyzer using the Virulence Factor Database (Miscellaneous)

| Isolate ↓ | Virulence Factor Class | | | | |
|---|---|---|---|---|---|
| | Toxin | | Miscellaneous | | |
| | Virulence factors | | | | |
| | Colicin-like Usp | Hemolysin/cytolysin A | Enterotoxin | Fimbrial adherence determinants | Misc. |
| *E. coli* [EC958] | *usp* | *hlyE/clyA* | | | |
| *K. quasipneumoniae* | | | | | |
| *K. pneumoniae* | | | | | |
| *E. roggenkampii* | | | | *bcfA, bcfB, bcfC, bcfD, bcfF, bcfG, csgA, csgB, csgC, csgD,csgF, csgG, fimA, fimC, fimD, fimF, fimH, fimI, fimZ, stjB, stjC, cheB, cheR, cheW, cheY, cheZ, motA, stjB, stjC* | Endotoxin LOS: *htrB, lgtF*. Magnesium uptake: *mgtC,* Manganese uptake: *mntB.* Motility flagella: *motB.* Non-fimbrial adhereance: *misL, shdA.* Biofilm formation: *adeG, pgaC.* Acid resistance: *ureB, ureG.* |
| *E. hormaechei* [19Y000094] | | | *ast,* | *fimA, fimC, fimD, fimF, fimH, fimI, fimZ, stjB, stiB, stkA, stkB, stkC, stkD, stkE, stkF, csgA, csgB, csgC, csgD, csgE, csgF, csgG, bcfC, lpfC,* | Biofilm formation: *adeG, pgaC.* Endotoxin LOS: *htrB, lgtF.* Motility, flagella: *flgB, flgC, flgD, flgE, flgF, flgG, flgH, flgI, flgJflgK, flgL, flgM, flhA, flhB, flhC, flhD, fliA, fliE, fliF, fliG, fliH, fliI, fliJ, fliM, fliN, fliP, fliQ, fliR, fliS, fliZ, flaA, motB.* Immune evasion: *gtrB.* Manganese uptake: *mntB.* Acid resistance: *ureB, ureG.* |
| *E. hormaechei* [19Y000373] | | | | *fimA, fimC, fimD, fimF, fimH, fimI, fimZ, stgA, stgB, stiB, orf01081, orf02252, stjB, stjC, stkA, stkC, stkD, stkE, stkF, csgA, csgB, csgC, csgD, csgE, csgF, csgG, bcfC, lpfC,* | Biofilm formation: *adeG, pgaC.* Endotoxin LOS: *htrB, lgtF.* Immune evasion: *galE, rmlA.* Manganese uptake: *mntB.* Trehalose-recycling ABC transporter: *sugC.* Acid resistance: *ureB, ureG.* O antigen: *orf01793; orf04415; orf04416; orf04419; orf04420.* Stress adaptation: *katG, sodC.* |
| *E. coli* [20Y000092] | *usp* | *hlyE/clyA* | *senB* | *lpfB, lpfC, lpfE* | Non-LEE encoded TTSS effectors: *espL1* |

**Table S3E** Virulence factors predicted by VFanalyzer using the Virulence Factor Database (Miscellaneous)

| Isolate ↓ | Virulence Factor Class - Miscellaneous | | | |
|---|---|---|---|---|
| | Autotransporters | Invasion | Antiphagocytosis | Serum Resistance |
| | Virulence factors | | | |
| | | | Capsule | LPS rfb locus |
| *E. coli* EC958 | *tibA, aida, aatA, agn43, cah, cdiA, cdiB, ehaA, ehaB, air/eaeX, espC, espI, espP, pet, pic, sat, tsh, upaG/ehaG, upaH, vat* | *ibeA, ibeB, ibeC, tia* | | |
| *K. quasipneumoniae* | | | *orf00393; orf00394; orf00395; orf00396; orf00397; orf00398; orf00407; orf00409; orf00410; orf00411; orf00412; orf00413; orf00416; orf00417; orf03577* | *orf00418; orf00419; orf00420; orf00421; orf00422; orf00423* |
| *K. pneumoniae* | | | *orf00780; orf02558; orf02559; orf02560; orf02561; orf02563; orf02564; orf02565; orf02573; orf02574; orf02575; orf02576* | *orf02579; orf02580; orf02581; orf02582; orf02583; orf02584; orf02585* |
| *E. roggenkampii* | *cdiB, ehaB* | *ibeB* | *orf02680; orf02688; orf02689; orf02695; orf02696; orf02697; orf02699; orf02700; orf02701; orf02708; orf02710; orf02743; orf02745, galF, ugd, wcaI, wzb* | *orf02687, misL, rmlD,* |
| *E. hormaechei* 19Y000094 | *ehaB* | *ibeB, cheB, cheR, cheW, cheY, cheZ, motA* | *orf01825, orf01829, orf01830, orf01834, orf01839, orf01840, orf01841, orf01842, orf01844, orf0184, orf01852, orf01853, orf01854, orf01885, orf01887, orf02058, orf02059, wbfY.* | *orf01826* |
| *E. hormaechei* 19Y000373 | *ehaB* | *ibeB, cheB, cheR, cheW, cheY, cheZ, motA, motB* | *wzb, orf00016, orf01745, orf02470, orf00018; orf01744; wzc; ugd, orf04397; orf04398; orf04408; orf04414; orf04415; orf04416; wcaI, orf04419; orf04420; orf04427; orf04428; orf04429* | *orf04394, orf04396* |
| *E. coli* 20Y000092 | *ehaB, upaG/ehaG, upaH, vat* | *ibeA, ibeB, ibeC* | | |

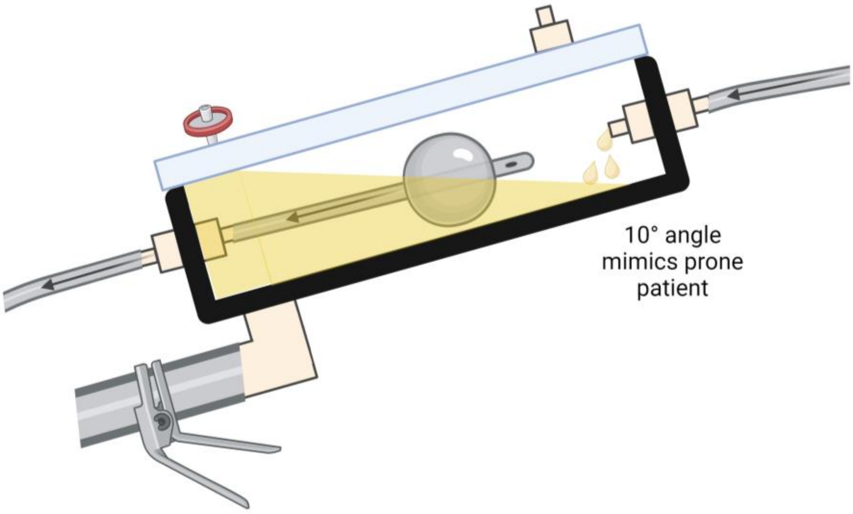


**Fig. S1** Schematic diagram of the Drip Flow Reactor® Foley catheter contamination model

The schematic depicts the cross section of a channel of the Drip Flow Reactor® when set up to run as the Foley catheter contamination model. The catheter is contaminated by dipping into a cell suspension prior to fixing within the chamber. Artificial urine medium is dripped into the chamber via one peristaltic pump and drains out of the chamber by gravity. The effluent tube at the base of the chamber allows for testing of the standing urine within the chamber.

**Fig. S2** CAUTI isolates plated onto UTI ChromoSelect agar

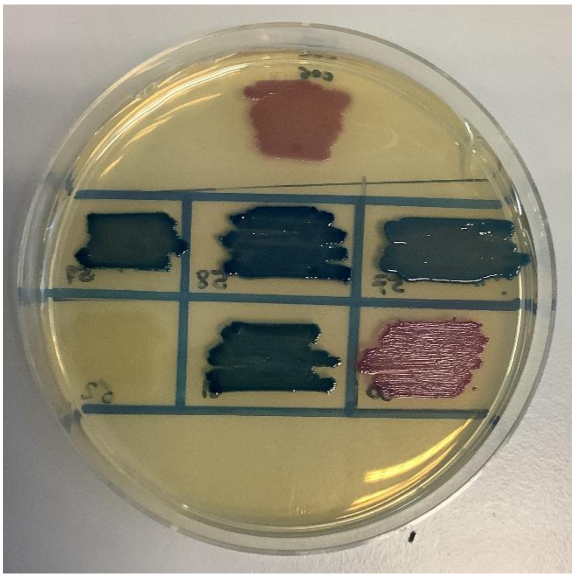

Strains plated out onto UTI ChromoSelect agar as per manufacturer's guidance and incubated at 37°C overnight. Top row: *E. coli* EC958, middle row (left to right): *E. roggenkampii*$^{18Y001733}$, *K. pneumoniae*$^{18Y001710}$, *K. quasipneumoniae*$^{18Y000138}$, bottom row (left to right): *E. coli*$^{20Y000092}$, *E. hormaechei*$^{19Y000373}$, *E. hormaechei*$^{19Y000094}$.

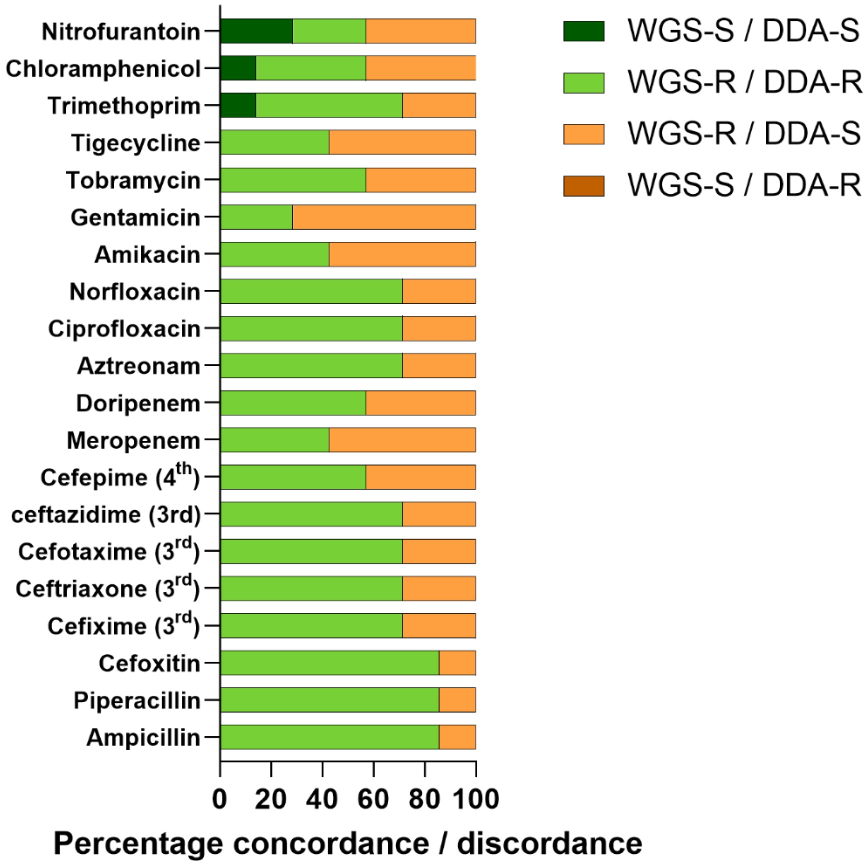

**Fig. S3** Concordance and discordance between genotypic and phenotypic datasets of antimicrobial resistance for seven (CA)UTI isolates.

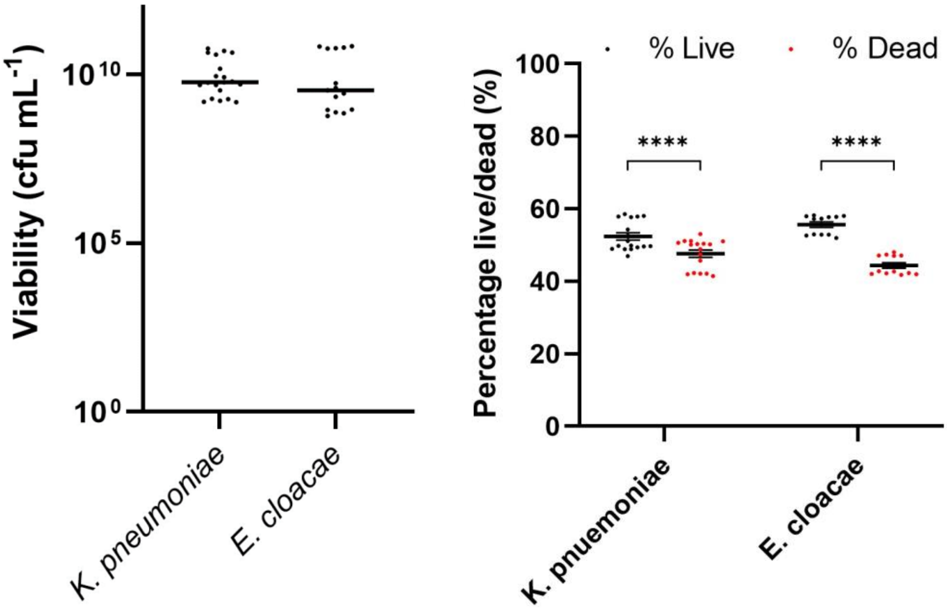


**Fig. S4** Viability of different genera of CAUTI causing pathogens

Proportions of live cells within five day old biofilms formed within the DFR biofilm contamination model were quantified by (A) determination of viable counts and (B) live/dead staining. N=3 ± SD.
